# Identifying novel subtypes of irritability using a developmental genetic approach

**DOI:** 10.1101/433342

**Authors:** Lucy Riglin, Olga Eyre, Ajay K Thapar, Argyris Stringaris, Ellen Leibenluft, Daniel Pine, Kate Tilling, George Davey Smith, Michael C O’Donovan, Anita Thapar

## Abstract

**Objective:** Irritability is a common reason for referral to services, strongly associated with impairment and negative outcomes, but is a nosological and treatment challenge. A major issue is how irritability should be conceptualized. This study used a developmental approach to test the hypothesis that there are several forms of irritability, including a ‘neurodevelopmental/ADHD-like’ subtype with onset in childhood and a ‘depression/mood’ subtype with onset in adolescence.

**Method:** Data were analyzed in the Avon Longitudinal Study of Parents and Children, a prospective UK population-based cohort. Irritability trajectory-classes were estimated for 7924 individuals with data at multiple time-points across childhood and adolescence (4 possible time-points from approximately ages 7 to 15 years). Psychiatric diagnoses were assessed at approximately ages 7 and 15 years. Psychiatric genetic risk was indexed by polygenic risk scores (PRS) for attention-deficit/hyperactivity disorder (ADHD) and major depressive disorder (MDD) derived using large genome-wide association study results.

**Results:** Five irritability trajectory classes were identified: low (81.2%), decreasing (5.6%), increasing (5.5%), late-childhood limited (5.2%) and high-persistent (2.4%). The early-onset, high-persistent trajectory was associated with male preponderance, childhood ADHD (OR=108.64 (57.45–204.41), p<0.001) and ADHD PRS (OR=1.31 (1.09–1.58), p=0.005); the adolescent-onset, increasing trajectory was associated with female preponderance, adolescent MDD (OR=5.14 (2.47–10.73), p<0.001) and MDD PRS (OR=1.20, (1.05–1.38), p=0.009). Both trajectory classes were associated with MDD diagnosis and ADHD genetic risk.

**Conclusions:** The developmental context of irritability may be important in its conceptualization: early-onset persistent irritability maybe more ‘neurodevelopmental/ADHD-like’ and later-onset irritability more ‘depression/mood-like’. This has implications for treatment as well as nosology.

## Introduction

Irritability – a heightened propensity to anger, relative to peers – is a common reason for referral to mental health services, is strongly associated with impairment and long-term adverse outcomes (1–6), yet remains a nosological and treatment challenge (1, 2). Currently, it is treated as a homogenous construct, but it is a core or accompanying feature of several psychiatric disorders and such differential associations suggest that subtyping may be necessary. This study set out to examine the possibility that there are multiple forms of irritability, including a ‘neurodevelopmental’ subtype with onset in childhood and a ‘depression/mood’ subtype with onset in adolescence.

Childhood irritability has typically been considered a feature of Oppositional Defiant Disorder (ODD) (7) - in the forthcoming ICD-11 it is likely to be considered a specifier of ODD. However irritability has been shown to be distinct from other ODD dimensions (headstrong, hurtful) in that it shows phenotypic and genetic associations with unipolar depression (5, 8). The DSM-5 has categorised severe, chronic childhood irritability as Disruptive Mood Dysregulation Disorder (DMDD) and grouped it with the mood disorders (9).

Yet irritability – and, more broadly, emotional dysregulation – is an especially common feature of attention-deficit/hyperactivity disorder (ADHD), which is grouped as a neurodevelopmental disorder under DSM-5. Irritability prevalence rates of 91% have been reported in children with the disorder (10). Evidence of clinical overlap between irritability and ADHD (11, 12), genetic overlap with ADHD, as well as features such as its manifestation in early development and male preponderance also have led recently to the suggestion that irritability should perhaps be conceptualised as a neurodevelopmental/ADHD-like problem, rather than a mood problem (12).

Two differentiating factors between neurodevelopmental and mood problems are developmental course and sex preponderance. Neurodevelopmental problems typically onset early, decline across childhood/adolescence and are more common in males, while mood problems tend to onset in adolescence and are more common in females (13). A developmental approach may therefore help towards better establishing whether irritability it is more appropriately conceptualised as a mood or neurodevelopmental problem. Indeed, a recent population-based study of irritability observed different developmental patterns for males and females: childhood irritability was more common in boys (for whom levels tended to decrease with age) while adolescent irritability was more common in girls (for whom levels tended to increase with age) (12). These findings raise the possibility that there may be different “subgroups” of irritability that have different developmental patterns and gender ratios - to our knowledge age-at-onset in childhood compared to adolescence has not previously been investigated as a possible source of heterogeneity in irritability.

The aim of this study was to take advantage of a longitudinal population-based cohort and use a developmental approach to test the hypothesis that there are at least two forms of irritability: one ‘neurodevelopmental/ADHD-like’ subtype with onset in childhood and one ‘depression/mood’ subtype with onset in adolescence. Specifically, we used a latent growth modelling approach to test the following hypotheses suggested by this formulation of irritability: (a) an irritability trajectory defined by an early age-at-onset would be associated with male gender, ADHD genetic liability as indexed by ADHD genetic risk scores (polygenic risk scores: PRS) and a higher rate of diagnosis of ADHD in childhood, and (b) an irritability trajectory defined by an age-at-onset around early-mid adolescence would be associated with female gender, depression genetic liability as indexed by major depressive disorder (MDD) genetic risk scores, and a diagnosis of MDD in adolescence.

## Method

### Sample

The Avon Longitudinal Study of Parents and Children (ALSPAC) is a well-established prospective, longitudinal birth cohort study. The enrolled core sample consisted of 14,541 mothers living in Avon, England, who had expected delivery dates of between 1^st^ April 1991 and 31^st^ December 1992. Of these pregnancies 13,988 children were alive at 1 year. When the oldest children were approximately 7 years of age, the sample was augmented with eligible cases who had not joined the study originally, resulting in enrollment of 713 additional children. The resulting total sample size of children alive at 1 year was N=14,701. Genotype data were available for 8,365 children following quality control. Ethical approval for the study was obtained from the ALSPAC Ethics and Law Committee and the Local Research Ethics Committees. Full details of the study, measures and sample can be found elsewhere (14, 15). Please note that the study website contains details of all the data that is available through a fully searchable data dictionary (http://www.bris.ac.uk/alspac/researchers/data-access/data-dictionary). Where families included multiple births, we included the oldest sibling.

Analyses were conducted including participants with at least two time-points of irritability data (see below) (N=7924): further details of sample sizes for available data are shown in Supplementary Figure 1.

### Irritability

In-line with previous work (7) irritability was defined using parent-reported data from the Development and Well-Being Assessment (DAWBA)(16) - a structured research diagnostic interview - at ages 7 years 7 months, 10 years 8 months, 13 years 10 months and 15 years 6 months. Three irritability items (severe temper tantrums, touchy and easily annoyed, angry and resentful) rated on a 3-point scale were summed to give a total score (0–6).

### Diagnoses

The DAWBA (16) were also used to assess ADHD, oppositional defiant disorder (ODD), conduct disorder (CD), general anxiety disorder (GAD) and major depressive disorder (MDD). Parent-reports were used in childhood (age 7 years 7 months) and to assess ADHD, ODD and CD in adolescence (15 years 6 months); self-report data were used to assess GAD and MDD in adolescence. DSM-IV diagnoses were generated using computer generated diagnoses (17); no diagnoses were mutually exclusive.

### Genetic liability

Polygenic risk scores (PRS) were used to capture common variant genetic liability for two disorders - Major Depressive Disorder (MDD) and Attention Deficit Hyperactivity Disorder (ADHD). For each disorder, PRS were generated as the weighted mean number of disorder risk alleles in approximate linkage equilibrium (r-square<0.1), derived from imputed autosomal SNPs using PRSice (18) (N=5559 of those included in this study: see Supplementary Figure 1). Sensitivity analyses were conducted using inverse probability weighting (19) to assess the impact of missing data (see Supplementary Material). Risk alleles were defined as those associated with case-status from the latest genome-wide association studies of MDD (135,458 cases and 344,901 controls) (20) and ADHD (19,099 cases and 34,194 controls) (21). Primary analyses defined risk alleles as those associated at a threshold of p<0.05; associations across a range of p-thresholds are shown in Supplementary Figure 2. Scores were standardized using Z-score transformation. Genotyping details as well as full methods for generating the PRS are presented in the Supplementary Material.

### Analyses

Growth mixture modelling (GMM) was conducted to identify developmental trajectories of irritability across ages 7 to 16 years in Mplus (22). GMM aims to group individuals into categories (trajectories) based on patterns of change across multiple time-points, with individuals within each category assumed to have the same growth curve (23). Starting with a single *k*-class solution, *k*+1 solutions are fitted until the optimum solution is reached. Models were run using a robust maximum likelihood parameter estimator (MLR) and full information maximum likelihood (FIML) estimation (24). As recommended, the optimal number of categories was determined by interpretability as well as model fit indices (see Supplementary Material)(23). A bias-free three step approach was used to investigate associations with PRS, gender and diagnoses (manual 3-step) and to estimate/compare prevalence rates (DCAT), which accounts for measurement error in class assignment (25); multinomial odds ratios are reported. Inverse probability weighting (19) was used to assess the impact of missing genetic data, which weights observations based on measures assessed in pregnancy that were predictive of variables in the analysis and/or inclusion in the subsample with genetic data (see Supplementary Material). Sensitivity analyses were conducted including gender as a covariate.

## Results

### Irritability trajectories

We identified a five-class solution (see Supplementary Material), characterised by distinct irritability trajectory classes: low (81.2%), decreasing (5.6%), increasing (5.5%), late-childhood limited (5.2%) and high-persistent (2.4%), shown in Figure 1.

**Figure 1.**
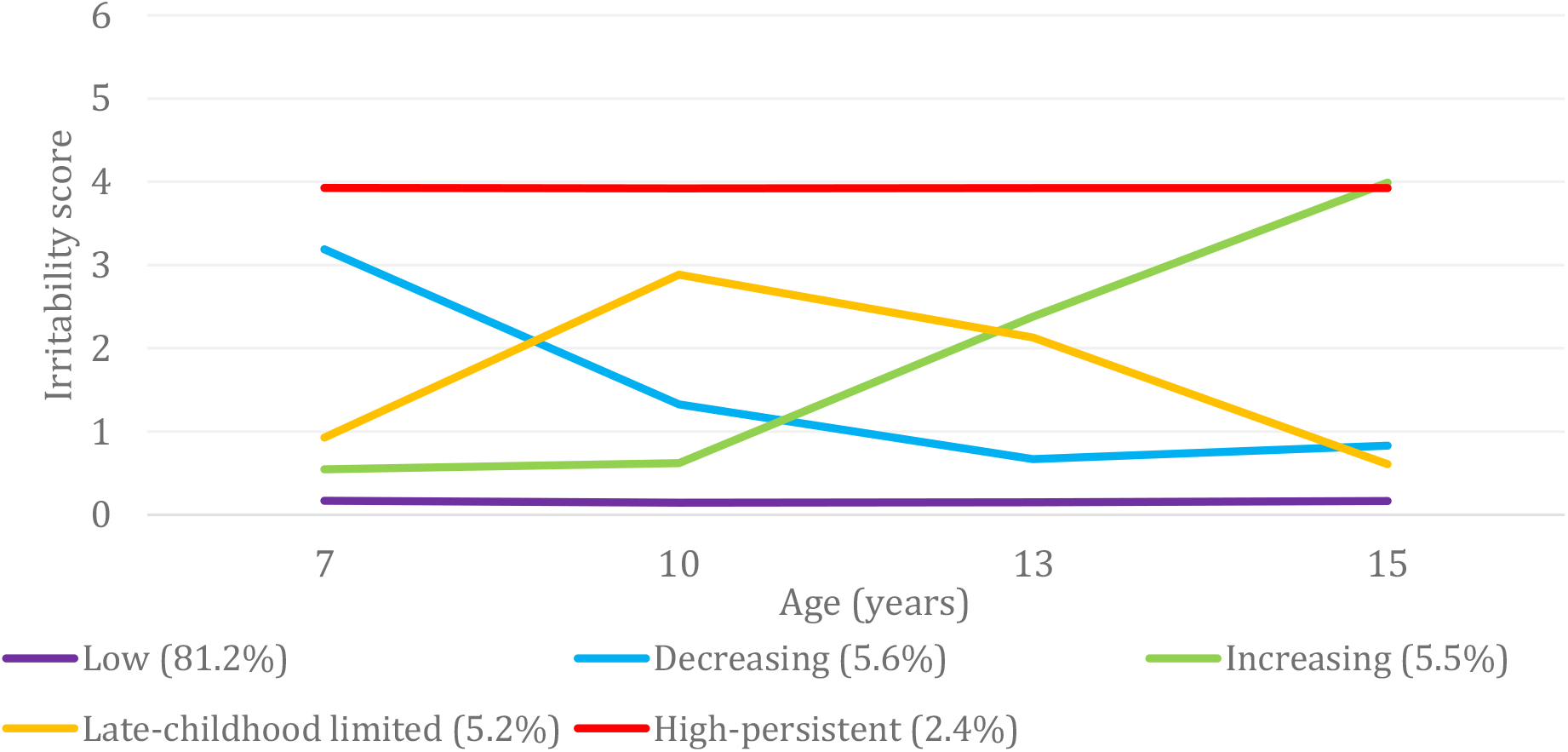
Irritability trajectories by class

A male preponderance was observed for the decreasing, late-childhood limited and high-persistent trajectory classes (55.7%, 55.7% and 63.7% male) and a female preponderance for the increasing trajectory class (40.5% male). Accordingly, male gender was associated with an increased likelihood of being in the decreasing (OR=1.27(1.01–1.59), p=0.038), late-childhood limited (OR=1.37 (1.09–1.73), p=0.007) or high-persistent classes (OR=1.76 (1.30–2.40), p<0.001) and a decreased likelihood of being in the increasing trajectory (OR=0.68 (0.54–0.87), p=0.002) compared to the low trajectory class (49.8% male).

### Genetic risk

Associations between irritability trajectory classes and ADHD PRS and MDD PRS are shown in Table 1. Mean PRS for each of the trajectory classes are shown in Figure 2. Compared to the low trajectory class, ADHD PRS were associated with an increased likelihood of being in both the high-persistent (OR=1.31 (95% CI 1.09–1.58), p=0.005) and increasing (OR=1.28 (95% CI 1.11–1.48), p=0.001) trajectory classes, with a similar risk of being in either trajectory (high-persistent vs. increasing OR=1.02 (95% CI 0.81–1.29), p=0.854).

**Figure 2.**
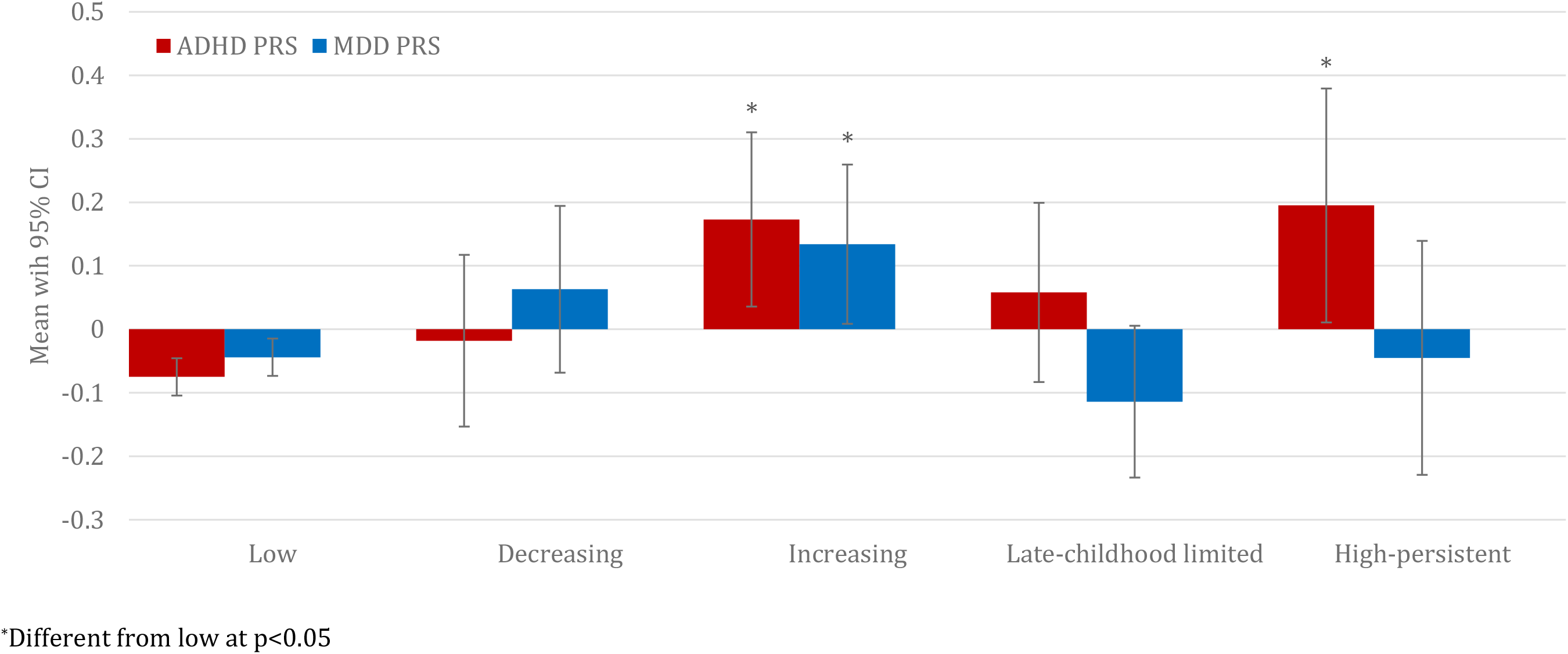
Mean ADHD and MDD genetic risk score by irritability trajectories, with 95% confidence intervals

**Table 1.**
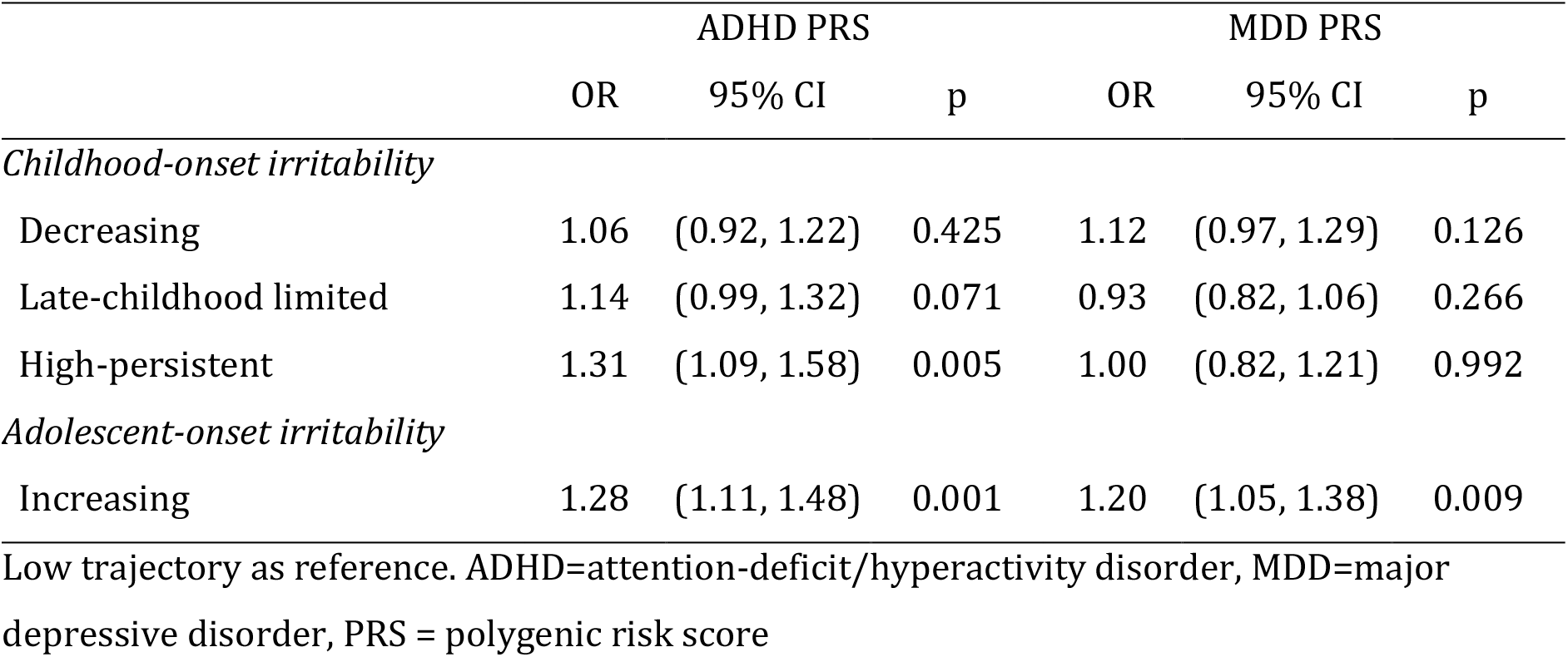
Association between ADHD and MDD genetic risk scores and irritability trajectories

MDD PRS were associated with an elevated likelihood of being in the increasing trajectory class compared to the low trajectory class (OR=1.20 (95% CI 1.05–1.38), p=0.009), although evidence for an elevated likelihood of being in the increasing trajectory class compared to the high-persistent trajectory class was weaker (OR=1.20 (95% CI 0.96–1.52), p=0.116).

Multivariable analyses including both PRS in the same model revealed the same pattern of results (see Table 1 vs. Supplementary Table 2) as did sensitivity analyses using inverse probability weighting to assess the impact of missing genetic data (Supplementary Table 3).

### Diagnoses

Estimated prevalence rates of ADHD, MDD, GAD, ODD and CD in childhood and adolescence are shown in Tables 2 and 3. Rates of all diagnoses varied across the irritability trajectory classes, with the exception that there was not strong evidence of variation in adolescent GAD across classes. At both developmental stages, rates of all diagnoses were generally highest in the high-persistent trajectory and were particularly high for ODD.

**Table 2.** Estimated prevalence of diagnoses in childhood across irritability trajectories

|  | ADHD<br>(N=7043) | MDD<br>(N=6947) | GAD<br>(N=7029) | ODD<br>(N=7034) | CD<br>(N=6979) |
| --- | --- | --- | --- | --- | --- |
| Low | 0.3% (0.1) | 0.1% (0.1) | 0.0% (0.0) | 0.1% (0.1) | 0.0% (0.0) |
| *Childhood onset irritability* |  |  |  |  |  |
| Decreasing | 9.3% (1.6) | 5.7% (1.2) | 1.1% (0.6) | 28.3% (2.5) | 4.0% (1.1) |
| Late-childhood limited | 6.3% (1.4) | 0.7% (0.5) | 0.5% (0.4) | 5.9% (1.5) | 1.0% (0.6) |
| High-persistent | 26.4% (3.4) | 4.2% (1.6) | 3.6% (1.4) | 51.0% (3.9) | 11.7% (2.5) |
| *Adolescent onset irritability* |  |  |  |  |  |
| Increasing | 1.9% (0.9) | 0.0% (0.0) | 0.0% (0.0) | 0.6% (0.7) | 0.2% (0.3) |
| $\chi^2(4)$ | 122.64<br>p<0.001 | 45.68<br>p<0.001 | 14.92<br>p=0.005 | 342.50<br>p<0.001 | 43.22<br>p<0.001 |

Standard errors in parentheses. ADHD=attention-deficit/hyperactivity disorder, MDD=major depressive disorder, GAD=generalised anxiety disorder, ODD=oppositional defiant disorder, CD=conduct disorder

**Table 3.** Estimated prevalence of diagnoses in adolescence across irritability trajectories

|  | ADHD<br>(N=4500) | MDD<br>(N=4900) | GAD<br>(N=4896) | ODD<br>(N=4490) | CD<br>(N=4489) |
| --- | --- | --- | --- | --- | --- |
| Low | 0.2% (0.1) | 1.1% (0.1) | 0.4% (0.1) | 0.1% (0.1) | 0.1% (0.1) |
| *Childhood onset irritability* |  |  |  |  |  |
| Decreasing | 1.7% (0.9) | 2.5% (1.1) | 0.8% (0.6) | 2.1% (1.2) | 1.1% (0.8) |
| Late-childhood limited | 0.6% (0.8) | 2.1% (1.1) | 0.8% (0.7) | 1.1% (1.5) | 0.7% (1.0) |
| High-persistent | 14.2% (3.4) | 7.3% (2.6) | 3.1% (1.8) | 60.9% (5.1) | 17.4% (3.8) |
| *Adolescent onset irritability* |  |  |  |  |  |
| Increasing | 7.3% (1.9) | 5.3% (1.6) | 2.9% (1.2) | 46.0% (3.9) | 17.0% (2.7) |
| $\chi^2(4)$ | 43.83<br>p<0.001 | 15.34<br>p<0.001 | 7.51<br>p=0.111 | 325.23<br>p<0.001 | 76.93<br>p<0.001 |

Standard errors in parentheses. ADHD=attention-deficit/hyperactivity disorder, MDD=major depressive disorder, GAD=generalised anxiety disorder, ODD=oppositional defiant disorder, CD=conduct disorder

#### Childhood ADHD

Childhood ADHD was associated with an increased likelihood of being in the decreasing (OR=30.97 (15.47–61.98), p<0.001) increasing (OR=5.89 (1.96–17.73), p=0.002), late-childhood limited (OR=20.39 (9.78–42.52) p<0.001) and the high-persistent (OR=108.64 (57.45–204.41), p<0.001) compared to the low trajectory class.

Comparing these four irritability trajectories, childhood ADHD was associated with a decreased likelihood of being in the increasing trajectory class (compared to decreasing OR=50.19 (0.07–0.52), p=0.001; late-childhood limited OR=0.29 (0.10–0.84), p=0.023; high-persistent OR=0.05 (0.02–0.15), p<0.001) and with the greatest likelihood associated with the high-persistent classes (compared to decreasing OR=3.50 (2.09–5.87), p<0.001; late-childhood limited OR=5.33 (2.92–9.73), p<0.001).

#### Adolescent MDD

Compared to the low trajectory class, an increased likelihood of adolescent MDD was found for the increasing (OR=5.14 (2.47–10.73), p<0.001) and high-persistent (OR=7.18 (3.10–16.61), p<0.001) trajectory classes (decreasing class OR=2.32 (0.88–6.12), p=0.088; late-childhood limited class OR=1.95 (0.63–6.04), p=0.250). Likelihood of adolescent MDD was similar for the high-persistent compared to the increasing trajectory class (OR=1.40 (0.51–3.84), p=0.516).

### Sensitivity analyses

Controlling for gender revealed the same pattern of results for both PRS and diagnoses (Supplementary Tables 4 and 5); mean PRS and estimated prevalence rates for diagnoses by gender are shown in Supplementary Figures 3 and 4).

## Discussion

This study aimed to investigate developmental trajectories of irritability across childhood and adolescence to test the hypothesis that there are different forms of ‘neurodevelopmental’ and ‘depression/mood’ irritability. Specifically we hypothesised that a ‘neurodevelopmental/ADHD-like’ irritability trajectory would be defined by an early age-at-onset, a male preponderance and would be associated with an increased genetic liability to ADHD and ADHD diagnosis in childhood, whereas a ‘depression/mood’ irritability trajectory would be defined by a later age-at-onset, a female preponderance, and would be associated with increased genetic liability to depression and MDD diagnosis in adolescence.

We identified five distinct developmental trajectory classes of irritability across childhood and adolescence. Four classes were characterized by elevated levels of irritability during at least some of this developmental period. Two irritability trajectory classes were early-onset: one was defined by symptoms that decreased over time and the other by high symptoms that persisted (5.6% and 2.4% of the sample respectively). These two groups show very similar developmental patterns to ADHD; with some individuals showing persistence over time and others remitting. An additional unexpected trajectory with irritability onset in late childhood was defined by an increase in symptoms at around age 10 years and a subsequent decrease at around age 13 years (5.2% of the sample). That is the age of high school transition in the UK as well as the onset of puberty for many but we can only speculate as to the underlying mechanisms for this class as this group has not been previously described. The final trajectory was defined by increasing symptoms that showed a later onset, around adolescence (5.5% of the sample).

In line with our first hypothesis, the two irritability trajectories with early-onset (decreasing and high-persistent) were both associated with male gender and ADHD diagnosis in childhood, as was the late-childhood onset (late-childhood limited) class. The high-persistent trajectory was also associated with increased ADHD genetic risk scores - although the childhood-onset trajectories that did not have persistent symptoms (decreasing and late-childhood limited) were not. This is similar to findings on the developmental patterns of ADHD symptoms, that those with persistent compared to childhood-limited symptoms of ADHD have an increased genetic liability to ADHD (26). An association between irritability and ADHD PRS has been observed previously in the total sample as well as a clinical sample and accords with an earlier twin study that observed shared genetic links between ADHD and emotional lability (11, 12). Interestingly, the later-onset (increasing) irritability trajectory was also associated with ADHD PRS – although rates of childhood ADHD diagnosis were low. It may be that the phenotypic expression of ADHD genetic liability in this predominantly female group manifests as mood problems (see 27), although further work would be needed to investigate this. Our findings therefore support the suggestion of a neurodevelopmental/ADHD-like ‘type’ of irritability, which onsets early, has a male preponderance and is associated with ADHD.

In line with our second hypothesis, the trajectory with irritability onset in adolescence (the increasing trajectory class) was associated with female gender. This ‘depression/mood-type’ irritability trajectory class was also associated with depression genetic risk scores and MDD diagnosis in adolescence. This class was also associated with ADHD genetic risk scores. Previous analyses of this same sample had failed to observe association between irritability with MDD genetic risk scores although previous twin work had indicated genetic overlap between irritability and depression (8, 28). This had raised some doubts about the primary classification of severe irritability as a mood problem and raised the possibility that disruptive mood dysregulation disorder might be better regarded as a neurodevelopmental. Indeed it appears that ICD-11 is going to take a different approach to DSM-5 and conceptualize irritability as a specifier of ODD, and not include severe irritability / disruptive mood dysregulation disorder as a mood disorder (the approach taken by DSM-5). Our findings suggest that the association between irritability and depression genetic risk scores may be specific to a subtype of irritability that onsets during adolescence.

Regardless of whether the DSM-5 or ICD-11 stance to dealing with severe irritability is most valid, neither have taken a developmental approach. Our findings suggest that development matters. This view is not new (see 29) and has been applied to other phenotypes including antisocial behaviour (30) but perhaps has been forgotten because it poses substantial challenges to clinicians and researchers. For example, R-DoC would provide a helpful research framework for conceptualizing irritability dimensionally across multiple levels. Yet this framework, as well as those in DSM-5 and ICD-11, is not developmentally informed in that irritability is classified the same regardless of age (and age-at-onset). Our findings cast doubt on the validity of this approach.

In terms of how the trajectory classes relate to psychiatric diagnoses, diagnostic rates are relatively low because this is a population-based cohort; nevertheless, there are some notable observations. Interestingly, while the high-persistent ‘neurodevelopmental/ADHD-like’ irritability trajectory class was not associated with MDD PRS, this class showed a similar risk of adolescent MDD as the increasing ‘depression/mood-like’ irritability trajectory. Thus both trajectory classes are associated with risk of major depression in adolescence, although the mechanisms of this associations are likely different, for example increased risk for MDD in the early-onset irritability subtype may be driven by environmental factors, such as increased life events associated with irritability, rather than genetic risk for MDD - although further work into this would be needed. It is established that child neurodevelopmental disorders such as ADHD, as well as oppositional defiant disorder and conduct problems, are risk factors for later depression so it is perhaps unsurprising that the ‘neurodevelopmental’ irritability trajectory is also associated with depression, although emerging research suggests that the presence of irritability in those with ADHD confers additional risk of depression (31).

Consistent with previous work (2) we found elevated rates of other psychiatric disorders in those with elevated irritability. Unsurprisingly, given that irritability is a feature of ODD, rates of ODD were particularly high in the irritability trajectory classes. Rates of ODD followed a similar pattern to irritability levels, i.e. in childhood, rates of diagnosis were highest in the childhood-onset irritability trajectories and in adolescence, were highest in the later-onset and high-persistent trajectories. This is somewhat unsurprising given that irritability is a core component of ODD and irritability was measured by ODD items - although a large proportion did not have ODD, suggesting that irritability in those without ODD is also important and informative. Interestingly we did not find strong evidence of variation in rates of adolescent GAD across the trajectories. Previous research on links between irritability and depression has often included anxiety symptoms in the same measure (28), or found the same pattern of results for depression and anxiety (6). Our findings suggest specificity to depression.

Our findings should be considered in light of some limitations. First, ALSPAC is a longitudinal birth cohort study that suffers from non-random attrition and those with increased genetic liability to disorder and with higher levels of psychopathology are more likely to drop out of the study (32, 33). However, our trajectory analyses used full information maximum likelihood (FIML) estimation (24) so that complete data on irritability were not required and inverse probability weighted (IPW) analyses suggested that missingness of genetic data did not have a large effect on our findings. Also, while PRS are useful indicators of genetic liability, ADHD and MDD PRS currently explains a minority of common variant liability to the disorder (20, 21) and so should be regarded as indicators of genetic liability rather than as predictors. Further, we did not include covariates in our analyses of ADHD and MDD diagnoses as we were interested in describing observed associations: this means that we cannot infer any causal associations between irritability and diagnoses. Different methods such as Mendelian randomization (MR) would be needed to investigate such research questions (34). Finally, our aim was to use a developmental approach to investigate nosology. However diagnostic and subgroup overlap is the rule in Psychiatry and this study was no exception. Our ‘neurodevelopmental/ADHD-like’ and ‘depression/mood-like’ irritability classes were both associated with ADHD and MDD diagnoses and ADHD genetic risk scores. We observed different patterns for these classes rather than identifying completely distinct groups.

In conclusion, our study identified different developmental trajectories of irritability including one with characteristics typical of neurodevelopmental/ADHD-like problems – early-onset, male preponderance and clinical and genetic links with ADHD - and one with characteristics typical of depression/mood problems - later-onset, female preponderance and clinical and genetic links with MDD. Both groups were associated with risk of adolescent MDD; both were associated with ADHD genetic risk scores. Overall the findings suggest that the developmental context of irritability may be important in its conceptualization; that has implications for treatment as well as nosology (2).

## Disclosures and acknowledgments

All authors report no competing interests.

We acknowledge the members of the Psychiatric Genomics Consortium for the publicly available data used as the discovery samples in this article. We thank the research participants and employees of 23andMe, Inc. for their contribution to this study. We are extremely grateful to all the families who took part in this study, the midwives for their help in recruiting them, and the whole ALSPAC team, which includes interviewers, computer and laboratory technicians, clerical workers, research scientists, volunteers, managers, receptionists and nurses. The UK Medical Research Council and the Wellcome Trust (102215/2/13/2) and the University of Bristol provide core support for ALSPAC. GWAS data was generated by Sample Logistics and Genotyping Facilities at the Wellcome Trust Sanger Institute and LabCorp (Laboratory Corporation of America) using support from 23andMe. We thank Alexander Richards and Richard Anney for preparing the quality controlled genome-wide association study summary statistics. This study was supported by the Wellcome Trust (204895/Z/16/Z).

